# Modeling Dynamics of Contact Inhibition of Proliferation and Structural Order in a Confluent Epithelium

**DOI:** 10.64898/2026.08.26.747344

**Authors:** Jiya Ghosh, Tapomoy Bhattacharjee, Sayantan Dutta

## Abstract

Contact inhibition of proliferation (CIP) enables epithelial tissues to self-regulate growth and maintain tissue homeostasis. However, how cell-level mechanical contact, tissue-scale structural order, and proliferation kinetics interplay remains a fundamental open question in living matter physics. Here, we present a particle-based model of a confluent epithelial monolayer governed by overdamped dynamics, where individual cells interact via a two-dimensional hard core-soft shoulder potential. By comparing structural evolution during quasistatic densification with previously reported experimental division kinetics, we find that the dynamics of proliferation arrest mimics the onset of direct steric contacts between the hard cores of the shell. Identifying hard core contacts as the physical driver of CIP, we couple our mechanical model with a stochastic Monte Carlo division scheme in which the instantaneous division rate decreases to zero from an intrinsic value as the number of hard core contact increases to six from zero. We demonstrate that for high intrinsic division rates, the cellular densification outpaces mechanical relaxation. This kinetic mismatch drives premature hard-core contact formation, shifts the onset of jamming and contact inhibition to lower packing fractions, and induces increasingly disordered transient configurations before the tissue universally converges to a hexagonal close-packed limit. Our model’s predicted division kinetics and structural order evolution are consistent with epithelial monolayer experiments, both reported and our own. This minimal physical framework links single-cell steric contact mechanics directly to tissue-scale growth regulation and structural evolution.

## 1 Introduction

The arrest of activity in a dense, disordered assembly of particles as packing fraction increases is one of the oldest and most general problems in soft matter physics, spanning colloidal suspensions, granular media, and foams [1]. Living tissues provide a striking biological realization of this phenomenon: as a population of dividing cells grows more crowded, proliferation itself shuts down once cells make sustained physical contact with their neighbors, a process termed contact inhibition of proliferation (CIP) [2]. Unlike passive colloidal or granular jamming, however, the particles here are active agents that generate their own density increase through division, coupling a structural, density-driven arrest to the internal kinetics of the densification. This coupling between packing geometry and a density-regulated internal degree of freedom (the decision to divide) makes confluent proliferating tissue a distinctive test bed for the physics of soft materials, in addition to being of central importance to tissue growth, development, and homeostasis, where unregulated proliferation is a hallmark of cancer [3].

The molecular basis of CIP is well characterized: E-cadherin-mediated adhesion and Hippo-pathway signaling (via nuclear YAP/TAZ localization) link mechanical stress at cell-cell junctions to cell-cycle arrest [4, 5], while an independent, cell-intrinsic checkpoint arrests the G1-to-S transition once a cell falls below a critical size [6], a rule shown to persist under tissue-scale confinement [7]. However, For the purpose of this article, the details of these pathways matter less than what they collectively achieve: they translate local crowding into a lower division rate, irrespective of the specific sensing mechanism.

At the tissue scale, this crowding-proliferation coupling manifests as genuine jamming and glassy phenomenology [8]. Traction-force measurements in migrating epithelial sheets reveal the stress fields that accompany collective expansion [9], motivating continuum descriptions in which growth is governed directly by internal stress [10, 11]. Previous works show that a dense epithelial monolayers undergo a glass-like dynamical arrest [1], reductions in local cell area alone are sufficient to suppress division [12], and spatial confinement can arrest cell-cycle progression independent of biochemical signaling [13]. Theoretical treatments of describing an epithelial tissue have followed three main paradigms: particle-based models representing cells as discrete objects interacting via pairwise potentials [14–18]; vertex-based models describing epithelial mechanics through explicit cell junctions and polygonal geometry [19–23]; and continuum descriptions treating the tissue as a continuous medium [10, 24, 25]. Within these frameworks CIP is typically implemented as a density- or stress-dependent growth rule, whether through direct stress-growth coupling [10,11], pressure-clamp thresholds [26, 27], local-area-dependent division [12, 20], cell-intrinsic size thresholds [7], or cell-cycle oscillators coupled to area statistics [28].

Despite these earlier works, how micro-mechanical neighbor interactions and division kinetics in confluent tissues interact and influence each other remains an open question. In this article, we address this by focusing on contact inhibition of proliferation in a confluent epithelial monolayer, where cells fully occupy the tissue space while their number density progressively increases during densification. To model the mechanical rearrangement of cells, we employ a hard-core soft-shoulder interaction potential, originally developed to capture anomalous structural ordering dynamics arising from competing interaction length scales [16, 29–31]. By comparing the structural evolution of our system with experimentally observed kinetics of CIP, we propose a mechanism where each cell regulates its division rate based on its number of hard-core contacts with immediate neighbors. Finally, we systematically investigate how structural evolution depends on the ratio of the characteristic timescales of cell division and mechanical relaxation as well as demonstrate that our model predictions are consistent with experimental observations in confluent epithelial monolayers.

## 2 Model Description

We model a confluent epithelial monolayer as a two-dimensional system of interacting particles governed by athermal overdamped Langevin dynamics:

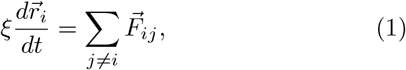

where 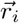 denotes the position of the *i*-th cell centroid, *ξ* is the effective translational drag coefficient, and 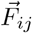 is the pairwise interaction force exerted on cell *i* by cell *j*.

To account for cell elasticity and volume exclusion, each cell is modeled as an isotropic structure consisting of an incompressible central core of fixed diameter *σ* surrounded by a soft, deformable shoulder of diameter *λσ* [Fig. 1A,B]. The hard core represents the non-compressible essential cellular volume (e.g., nucleus and major organelles), whereas the soft shoulder represents the surrounding deformable cytoplasm that permits local cell overlap and structural rearrangements. The size ratio *λ* dictates the relative extent of the soft shoulder.

**Figure 1:**
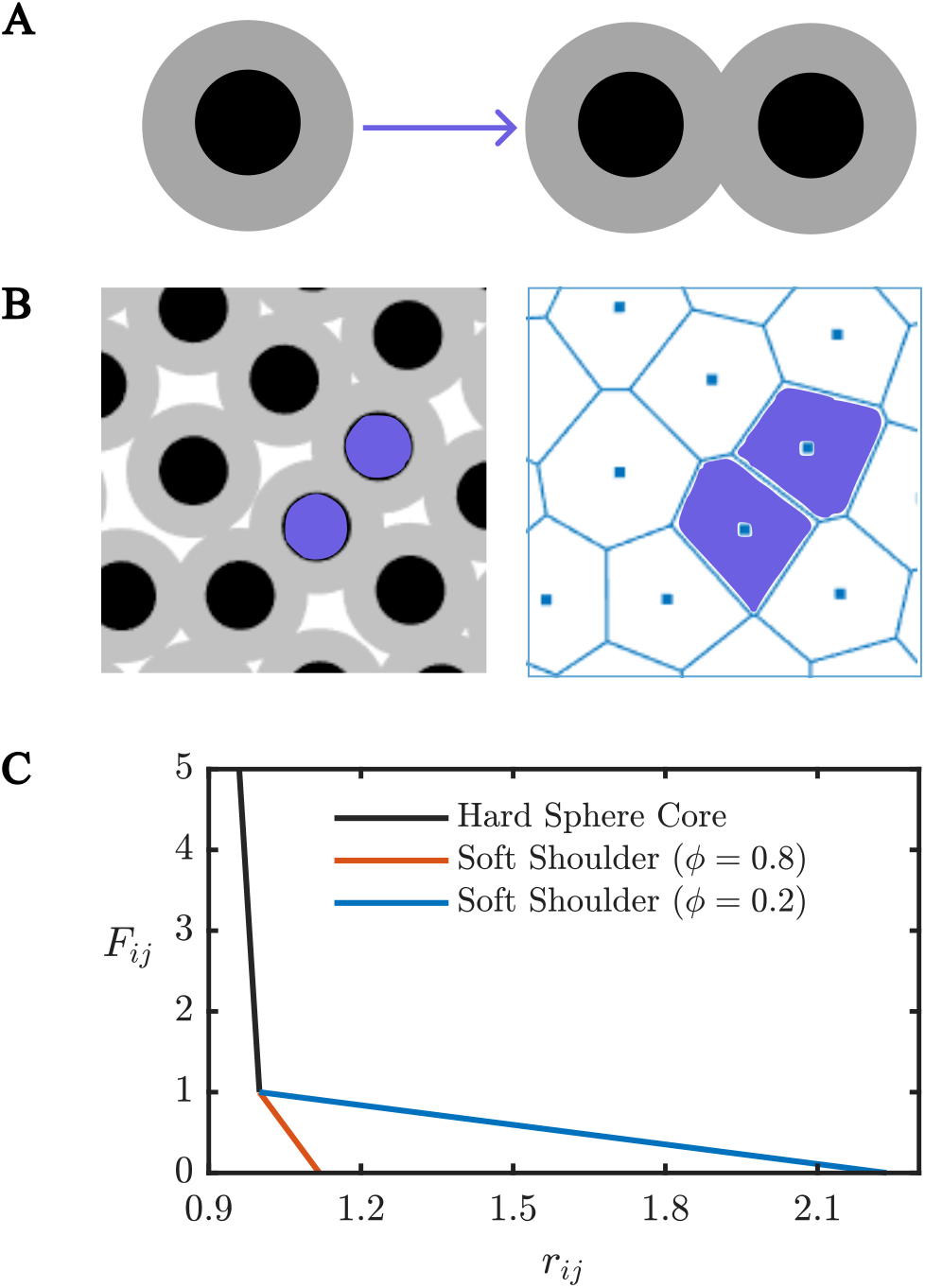
The process of cell division and hard core-soft shoulder interaction force-field: (A) Schematic of an individual cell modeled with a nearly incompressible hard core (black) and a soft shoulder (gray) undergoing division. The left and right hand side snapshots represent mother and daughter cells respectively. (B) Representative simulation snapshots of a tissue subregion following cell division (left), with the sister cell cores highlighted in purple, and the corresponding Voronoi tessellation based on cell centroids (right). Purple polygons indicate the Voronoi domains associated with the same cell pair following division. (C) Dimensionless interaction force 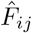 as a function of center-to-center distance 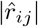. The steep black curve illustrates the stiff core repulsion 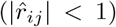, while blue and red curves represent soft-shoulder interactions 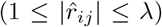 at high (*ϕ*_core_ = 0.8) and low (*ϕ*_core_ = 0.2) hard-core packing fractions, respectively.

We non-dimensionalize length by the hard-core diameter, 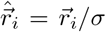, and time by the shoulder elastic relaxation timescale, 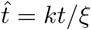, where *k* is the spring constant characterizing soft-shoulder elasticity. This yields the dimensionless equation of motion:

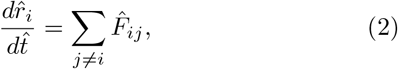

where the dimensionless interparticle force 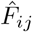 is defined by the piecewise function [Fig. 1C]

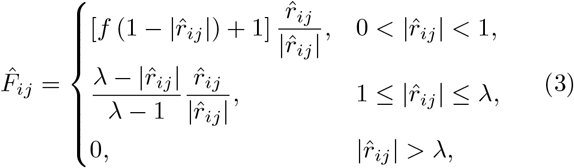

with *f* ≫ 1 representing the stiffness ratio between the hard core and the soft shoulder.

We initialize the system by placing cell centroids within a periodic square simulation domain with length *Lσ* using random sequential addition (RSA), ensuring that no hard-core overlaps occur. To represent initial confluency, the aggregate area of the soft shoulders is set equal to the box area, 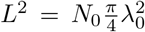, where *N*_0_ is the initial cell number and *λ*_0_ is the initial soft-shoulder aspect ratio. Cell division is implemented by splitting a mother cell into two daughter cells separated by a small, randomly oriented displacement vector. While the hard-core diameter *σ* remains invariant across generations, the available deformable volume within the domain is conserved and continuously redistributed among all cells [Fig. 1 A-C]. Consequently, as proliferation increases the total cell count (*N* → *N* + 1), maintaining domain area conservation 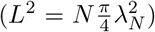 induces a dynamic rescaling of the soft-shoulder diameter:

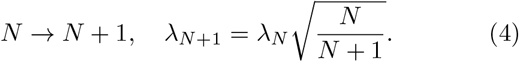

The spatial centroid positions are mapped to an explicit polygonal tissue morphology using a Voronoi tessellation [Fig. 1 B] to represent an epithelial tissue morphology.

## 3 Results

### 3.1 Quasistatic Division Dynamics and Contact Inhibition Mechanics

We begin our analysis by examining the quasistatic limit of cellular proliferation, where the timescale of division is vastly slower than the mechanical relaxation timescale of the tissue. The simulation domain is initialized at a hard-core packing fraction of 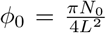. Proliferation is modeled by selecting a cell at random to undergo division, followed by a mechanical equilibration phase over a duration of 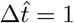. This cycle of division and equilibration is repeated iteratively until reaching a maximum hard-core packing fraction of *ϕ*_max_ = 0.9.

We characterize the structural and mechanical evolution during densification using two complementary metrics: (i) the average hard-core contact number per cell, *z*, (defined by pairwise distance criterion 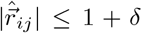, where *δ* ≪ 1 represents a small contact threshold), and (ii) the hexatic bond-orientational order parameter [23, 32],

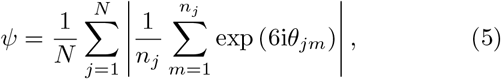

which quantifies global six-fold rotational symmetry. Here, *n*_*j*_ denotes the number of Voronoi neighbors of cell *j*, and *θ*_*jm*_ is the orientation angle of the bond vector connecting cell *j* to its *m*-th neighbor relative to a fixed reference axis.

By construction, *ψ* = 0 corresponds to an uncorrelated spatial distribution (Poisson), whereas *ψ* = 1 indicates a perfect triangular lattice.

Plotted against both average cell area (bottom axis) and hard-core packing fraction *ϕ* (top axis), these metrics reveal distinct structural behaviors during tissue crowding. The hard-core contact number *z* remains identically zero up to a critical packing fraction of *ϕ* ≈ 0.60, beyond which it increases sharply, approaching *z* ≈ 6 near close packing (Fig. 2A). In contrast, the hexatic order parameter *ψ* grows continuously throughout the densification process (Fig. 2B).

**Figure 2:**
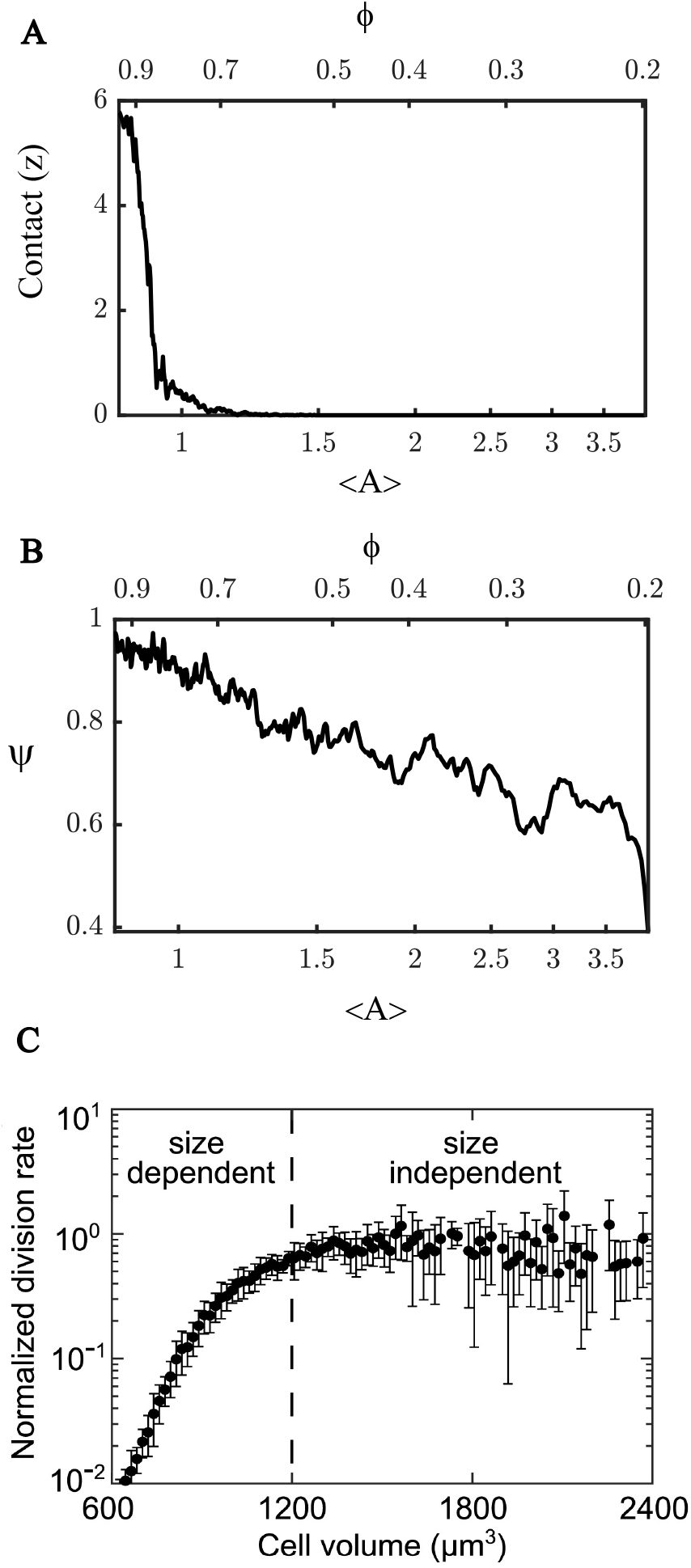
Dyanmics of structural order and contact mechanics under quasistatic cellular densification. Evolution of (A) the average hard-core contact number per cell, *z*, and (B) the hexatic bond-orientational order parameter, *ψ*, as functions of hard-core packing fraction *ϕ* (top axis) and mean cell area (bottom axis). (C) Experimentally observed division rate as a function of average cell area in an epithelial monolayer (Reproduced with permission from Devany et. al, 2023 [7]).

To elucidate the physical mechanism governing density-dependent proliferation arrest (contact inhibition), we compare the evolution of both *z* and *ψ* against experimentally measured cell division rates as a function of mean cell area in epithelial monolayers [7] (Fig. 2 C). Remarkably, the experimental division rate kinetics exhibit a sharp onset threshold that maps closely onto the contact number *z*, whereas they show poor correlation with the gradual rise of *ψ*.

These observations indicate that contact inhibition of proliferation in epithelial tissues may primarily be governed by physical proximity between stiff cellular cores (e.g., nuclei and organelle-dense regions), rather than by structured ordering or cell area variance as previously proposed in models of cell-cycle biochemical oscillators [28]. Mechanistically, we hypothesize that when incompressible cell cores are brought into close spatial contact, local steric compressive stresses trigger mechanotransductive signaling pathways that inhibit cell cycle progression.

### 3.2 Adaptive rate of Division based on contact

While the quasistatic framework captures the structural trajectory under densification, it does not accurately represent the realistic kinetics of cell proliferation. Specifically, the quasistatic protocol produces a linear increase in cell number over time, whereas growing tissues undergoing contact inhibition typically exhibit exponential growth at low densities, followed by progressive saturation as crowding takes effect.

Building upon our observation that proliferation kinetics are correlated with core-to-core contacts, we refine our mechanical framework to explicitly model the individual cell division rate as a function of local hard-core coordination [Fig. 3]. Specifically, we propose a contact-dependent division rate *K*_*i*_ for cell *i*:

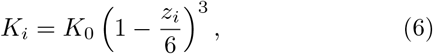

where *K*_0_ is the intrinsic division rate in the absence of hard-core contacts (*z*_*i*_ = 0), and *z*_*i*_ denotes the instantaneous hard-core coordination number of cell *i*. As local crowding causes the tissue to approach two-dimensional hexagonal close packing of the hard cores (*z*_*i*_ → 6), *K*_*i*_ vanishes continuously to zero, enforcing complete contact inhibition of proliferation.

**Figure 3:**
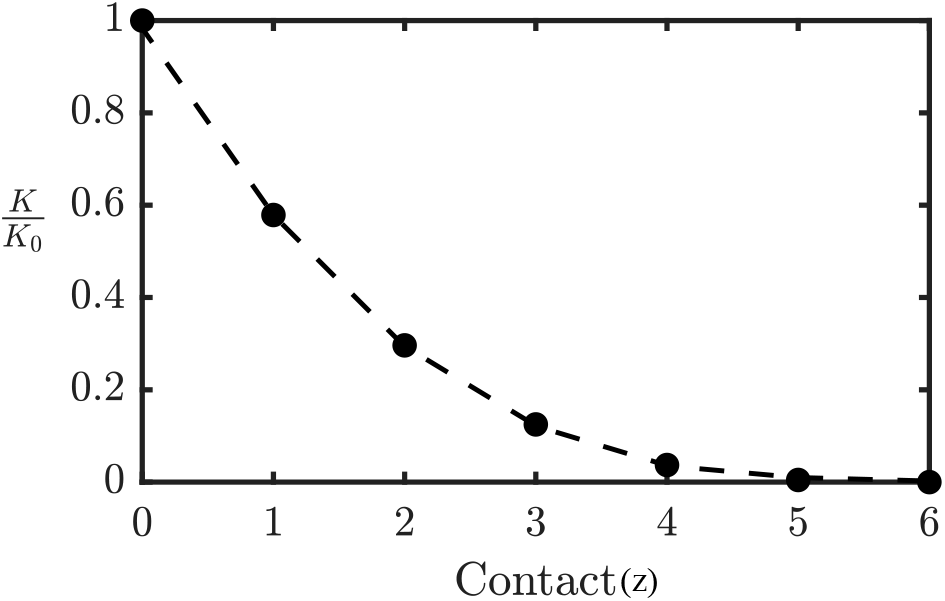
Contact-dependent division rate model and population growth kinetics. Normalized division rate *K/K*_0_ = (1 ™ *z/*6)^3^ as a function of the local hard-core contact number *z*.

We integrate this contact-dependent rate law into our mechanical framework by coupling the Langevin equations of motion with a stochastic Monte Carlo scheme. At each numerical integration timestep 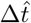, every cell evaluates its division probability as,

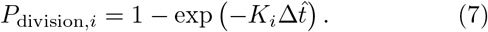

### 3.3 Division rate dependent structural evolution

Using the contact-dependent proliferation model, we investigate tissue densification dynamics across a range of intrinsic division rates *K*_0_, starting from a fixed initial packing fraction *ϕ*_0_ = 0.20 [Fig. 4 A,B]. As in the quasistatic case, we monitor the evolution of the hard-core contact number *z*, the hexatic bond-orientational order parameter *ψ* [Fig. 4 C,D], and the instantaneous division rate *K* [Fig. 4 E] as functions of the hard-core packing fraction *ϕ*. At maximum densification, the tissue universally approaches a hexagonal close-packed state (*z* → 6, *ψ* → 1) irrespective of *K*_0_. Crucially, however, the transient structural pathways leading to this fully ordered limit depend strongly on the division rate.

**Figure 4:**
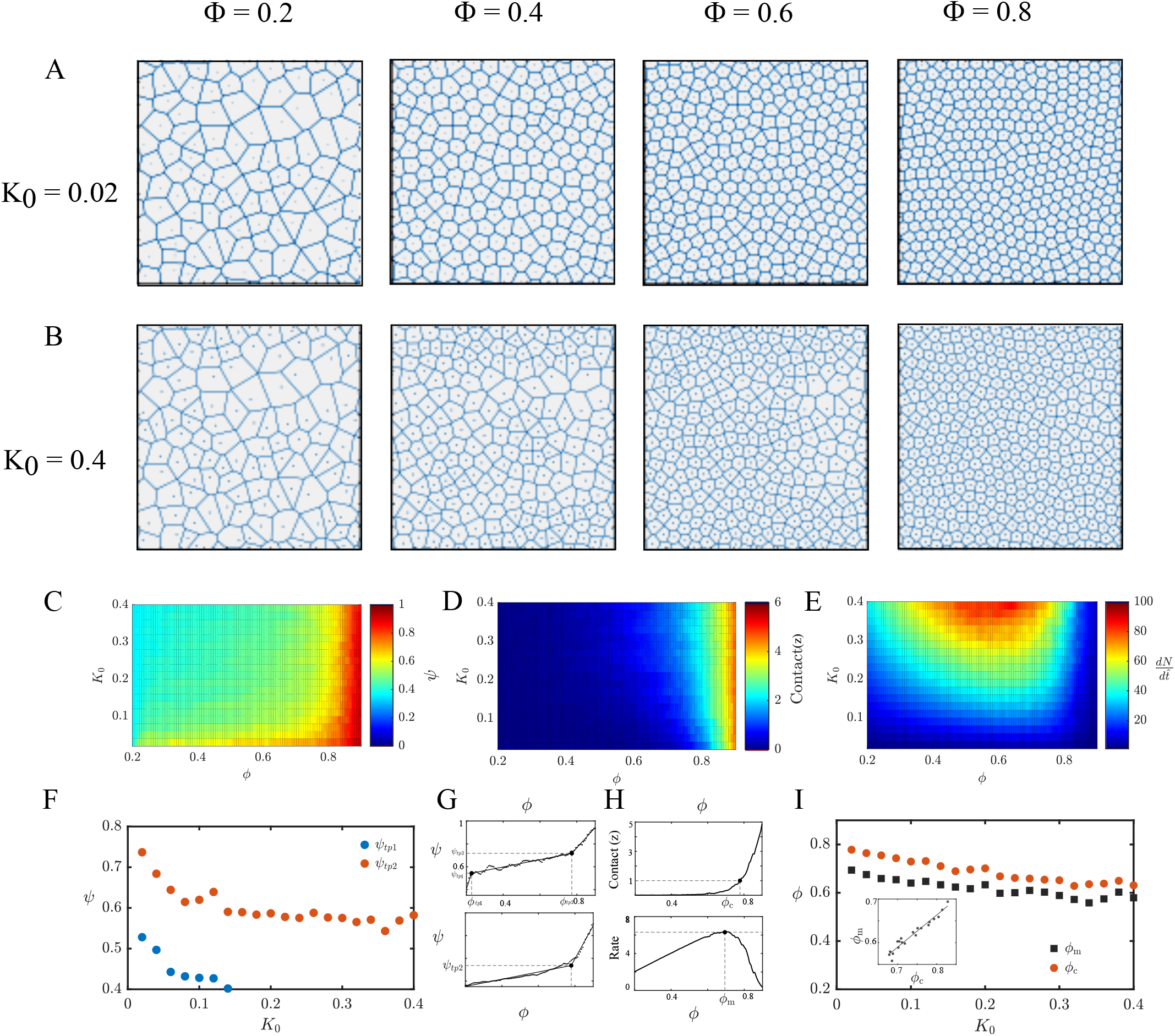
Rate-dependent structural ordering and contact mechanics: (A, B) Representative Voronoi tessellations of tissue morphology under (A) slow (*K*_0_ = 0.02) and (B) fast (*K*_0_ = 0.40) intrinsic division rates. (C) Hexatic bond-orientational order parameter *ψ*, (D) average hard-core contact number per cell *z*, and (E) instantaneous global proliferation rate 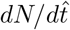 evaluated across varying packing fractions *ϕ* for different intrinsic division rates *K*_0_. (F) Order parameter values *ψ* at the structural transition points as a function of *K*_0_. (G) Representative piecewise linear fits demonstrating three-stage (top) and two-stage (bottom) ordering trajectories in *ψ* ™ *ϕ* space.(H) Methodological determination of the hard-core contact onset packing fraction *ϕ*_*c*_ (defined where *z* = 1, top) and the contact inhibition onset packing fraction *ϕ*_*m*_ (defined where 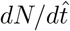 reaches its maximum, bottom). (I) Evolution of *ϕ*_*c*_ and *ϕ*_*m*_ as functions of intrinsic division rate *K*_0_. Inset: Scatter plot and a linear fit between *ϕ*_*c*_ and *ϕ*_*m*_.

Specifically, the hexatic order parameter *ψ* exhibits a distinct three-stage trajectory during densification: an initial rapid ordering phase, an intermediate slow ordering phase, and a final rapid ordering phase. These three regimes are well captured by piecewise linear fits with distinct slopes [Fig. 4 G]. Remarkably, the transition packing fractions separating these regimes are invariant with respect to *K*_0_, occurring at fixed thresholds of *ϕ*_tp1_ ≈ 0.23 and *ϕ*_tp2_ ≈ 0.78. In contrast, the order parameter values at these transition points decrease monotonically with increasing *K*_0_ [Fig. 4 F], demonstrating that faster proliferation generates increasingly disordered intermediate configurations [Fig. 4(A,B)]. Furthermore, while the slope *dψ/dϕ* in the intermediate slow ordering phase remains independent of *K*_0_, the slope in the initial fast ordering phase diminishes with increasing *K*_0_ and the phase eventually disappears for *K*_0_ *>* 0.15.

Summarily, these results indicate that tissue densification possesses an intrinsic structural relaxation trajectory characterized by a soft-shoulder-dominated slow ordering phase at moderate densities, followed by a hard-core-dominated fast ordering phase at high densities. In the initial phase, the system rapidly relaxes from the uncorrelated disorder of the random sequential addition (RSA) state toward this intrinsic trajectory. With increasing *K*_0_, the intrinsic path itself shifts toward higher structural disorder; consequently, the initial relaxation slope decreases until this stage disappears for *K*_0_ *>* 0.15. Conversely, in the final fast ordering regime, the slope steepens with increasing *K*_0_ up to *K*_0_ = 0.15. Because all systems must ultimately converge to the identical triangular close-packed limit (*ψ* = 1), tissues proliferating at higher rates are forced to undergo a more pronounced, rapid structural reorganization to recover from their highly disordered intermediate states.

Next, we examine the emergence of hard-core contacts and the resulting mechanical inhibition of proliferation. Consistent with the quasistatic regime, the tissue maintains zero hard-core contacts (*z* = 0) across a broad range of low-to-intermediate densities before undergoing a steep rise toward *z* = 6 at higher packing fractions [Fig. 4 D]. We quantify the onset of hard-core contact by identifying the characteristic packing fraction, *ϕ*_*c*_, where the average coordination number reaches *z* = 1 [Fig. 4 H]. Notably, *ϕ*_*c*_ shifts to systematically lower packing fractions as the intrinsic division rate *K*_0_ increases [Fig. 4 I], signifying a rate-dependent, pre-mature onset of structural jamming.

Concurrently, we analyze the kinetics of contact inhibition by tracking the global proliferation rate, 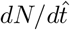, as a function of hard-core packing fraction *ϕ*. The global rate displays a characteristic non-monotonic profile across all *K*_0_: an initial rise driven by exponential population growth, followed by a rapid decay as steric contact inhibition takes effect. We define the onset of contact inhibition by the packing fraction *ϕ*_*m*_ at which the global division rate reaches its maximum [Fig. 4 H]. Paralleling the behavior of *ϕ*_*c*_, *ϕ*_*m*_ decreases monotonically with increasing *K*_0_ [Fig. 4 I] and exhibits a strong positive correlation with *ϕ*_*c*_ [Fig. 4 I, inset].

Collectively, these results reveal that rapid proliferation outpaces the structural relaxation timescale of the tissue. When *K*_0_ is high, cells lack sufficient time between division events to mechanically equilibrate and reorganize. This inability to relax drives premature steric contacts between stiff cellular cores (a hallmark of jamming) at lower overall densities, thereby triggering early contact inhibition and restricting tissue growth.

### 3.4 Validation against experimental monolayer dynamics

We next compare the dynamics predicted by our computational framework against epithelial monolayer experiments, both those reported previously and those we conducted ourselves. Specifically, we seeded MDCK (Madin–Darby Canine Kidney) cells, incubated under standard physiological condition (i.e. 37 ^*°*^ C, 5 % CO_2_), at approximately 70 % confluence onto a glass substrate and maintained in complete DMEM growth medium supplemented with 10% fetal bovine serum (FBS). For nuclear visualization, we stained the cells with Hoechst dye for 30 minutes. Finally we conducted time-lapse live-cell imaging using a confocal microscope in a 30 minute interval (Fig. 5A). The regime relevant to our model spans from the onset of confluence to the point at which the tissue begins to grow into the third dimension.

**Figure 5:**
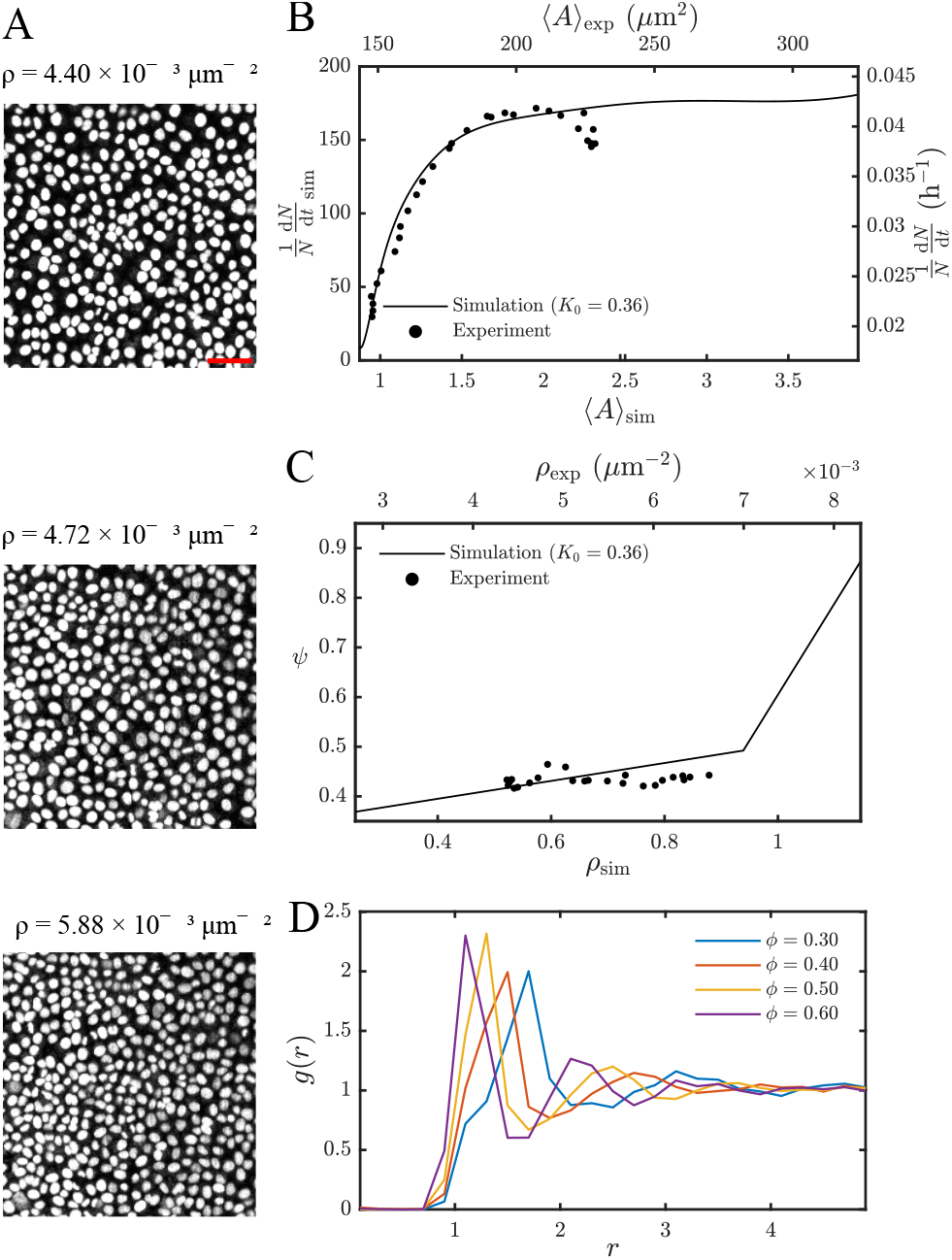
Comparison of simulation predictions to experimental outcome. (A) Time-lapse confocal microscopy snapshots of live Hoechst-stained MDCK cell monolayers at three representative densities (*ρ*) spanning the regime, where the tissue stays as confluent and as a monolayer. The scale bar in red denotes a length of 50 *µ*m. (B) Per capita division rate of cells as a function of mean cell area computed from simulations with adaptive rate of division (solid line, left and bottom axes) as well as from our experiment (scattered points, right and top axes). The experimental rate of division is computed from fitting a smooth curve to the observed *N*(*t*) and finding the derivative of the corresponding curve. (C) Hexatic bond order parameter as a function of number density computed from simulations with adaptive rate of division (solid line, bottom axis) as well as from our experiment (scattered points, top axis). (D) Radial distribution function *g*(*r*) obtained from model simulations across packing fractions *ϕ* = *{*0.3, 0.4, 0.5, 0.6*}* at an intrinsic division rate *K*_0_ = 0.2.

We first compare the per-capita division rate obtained from our adaptive simulation framework with that measured in our experiments. Both the simulation (by construction) and the experiment show the same qualitative trend: a constant per-capita division rate that begins to decrease once a threshold density is reached, consistent with the results of Devany *et al*. [7] (reproduced in Fig. 2C alongside our data in Fig. 5B). We next quantify structural order directly from the experimental images: after segmenting nuclei and extract-ing their centers at each density, we compute the bond-order parameter *ψ*. Its value fluctuates around 0.45 throughout densification, with the fluctuations remaining within the net increase in *ψ* at the slow-ordering regime (Fig. 5C).

Finally, we examine a structural signature characteristic of hard-core soft-shoulder systems in which the soft-shoulder radius is rescaled with density: the characteristic inter-particle spacing decreases with increasing density while the degree of structural (dis)order is preserved, until packing becomes tight enough for the hard core to dominate the interaction [16].. Our simulations with an adaptive division rate reproduce this behavior in the slow ordering regime — the position of the first peak of the radial distribution function (RDF) shifts systematically with density while its height remains nearly unchanged, and no long-range order emerges at any density within this regime (Fig. 5D). This is consistent with the earlier observation of Puliafito *et al*. [12] in a densifying confluent epithelial monolayer. Our own experiments show the same signature: the first RDF peak shifts in position between the early and lates stage of the experiment, while its height remains nearly the same [Supplementary Fig. S4]. Taken together, these results show that the slow-ordering phase captured by our simulations is consistent with the densification of an epithelial monolayer, both in its proliferation kinetics and in the dynamics of its structural order.

## 4 Conclusions

Summarily, we have introduced a particle-based framework featuring an adaptive core–shoulder interaction potential to model the mechanics and structural dynamics of confluent epithelial monolayers undergoing progressive densification [Fig. 1]. By simultaneously tracking the local hard-core contact number (*z*) and the global hexatic order parameter (*ψ*), we demonstrated that experimental proliferation arrest aligns with the sharp onset of hard-core steric contacts rather than the continuous rise of hexatic order [Fig. 2], suggesting steric core crowding as the primary driver of contact inhibition of proliferation (CIP). Encoding this mechanism into a contact-dependent division rate [Fig. 3], we showed that while the intrinsic division rate *K*_0_ leaves the terminal hexagonal close-packed state invariant (*z* → 6, *ψ* → 1), it fundamentally reshapes the transient structural path: higher *K*_0_ values shift the onset of steric contact (*ϕ*_*C*_) and peak division rate (*ϕ*_*m*_) to lower packing fractions, forcing the tissue through increasingly disordered intermediate configurations before late-stage rapid ordering restores the close-packed limit [Fig. 4]. We further validated this framework against epithelial monolayer experiments, both from this studies and the previous sones, showing that both the predicted division kinetics and the density-dependent evolution of structural order are consistent with experimental observations [Fig. 5].

The broader significance of this work rests on establishing a minimal, mechanistically transparent link between single-cell steric interactions and tissue-scale growth regulation. By demonstrating that CIP can be quantitatively captured through hard-core steric contacts alone, without invoking explicit biochemical oscillators or complex stress-signaling cascades, our model provides a physically intuitive and computationally efficient framework to study the dynamics of epithelial monolayer. Moreover, our work depicts a charateristic disorder to order transition dynamics during densification. Specifically, our findings highlight that the competition between cell division and mechanical relaxation timescales dictates the structural trajectory of a densifying tissue, establishing the proliferation rate itself as a tunable physical control parameter for tissue-scale order and structural jamming.

Several promising avenues remain for future exploration. From a biological perspective, identifying the precise subcellular components that constitute the incompressible “hard core”, such as the nucleus, dense organelle crowding, or the perinuclear actin network, presents an exciting challenge for experimental characterization. A separate limitation concerns the interpretation of “proliferation arrest” itself: in real monolayers, our experiments show that the cells do not often simply stop dividing once packing becomes tight. Instead, they relieve steric crowding by extruding or growing into the third dimension. On the other hand, extending this hard-core soft-shoulder formulation to three dimensions would test whether a similar contact-mediated mechanism governs growth arrest in inherently three-dimensional living systems, such as tumor spheroids and organoids, since the dynamics of structural order can differ markedly between two and three dimensions.

From the computational side, our current model treats cells as isotropic particles with no implicit fluctuations. Examining how shape anisotropy and active motility, representing cell migration, influence structural order dynamics would be of significant biological relevance. Similarly, explicitly coupling division orientation to local mechanical anisotropy could uncover the relation between proliferation kinetics and tissue-scale stress patterning. A complementary direction is to recast the same physical principle within a vertex-model framework: rather than an interparticle hard core, division could be suppressed by a steep energy penalty that switches on once a cell’s apical area falls below a critical threshold, allowing direct comparison between particle-based and vertex-based descriptions of the same crowding-induced arrest. Finally, the impact of cellular heterogeneity in size, stiffness, and proliferative capacity on the dynamics of structural order is another question worth exploring. Taken together, these extensions chart a path toward a unified, minimal physical description of how mechanics, packing, and division kinetics jointly sculpt the growth of living tissue.

## Author contributions

J.G. developed the simulation code, analyzed the result, performed the experiment and analyzed the experimental results. S.D. and T.B. conceptualized the project and acquired funding. S.D. and J.G. prepared the manuscript. All authors have read and edited the manuscript.

## Acknowledgements

We thank Sitabhra Sinha, Shakti Menon and Solingyur Zimik for helpful discussion. J.G. thanks Anushka Chowdhury for her help with the experimental setup. S.D. acknowledges PMECRGA grant from Anusandhan National Research Foundation (ANRF) vide file no ANRF/ECRG/2024/001067/ENS. TB and SD acknowledge the Department of Biotechnology, Government of India, Project Identification Number BT/PR54128/BMS/85/596/2024. TB acknowledges the support from Department of Atomic Energy, Government of India vide Project Identification No. RTI 4006, Nikon India, as well as the EMBO Young Investigator Program.

