## Supplementary Material for "Modeling Dynamics of Contact Inhibition of Proliferation and Structural Order in a Confluent Epithelium"

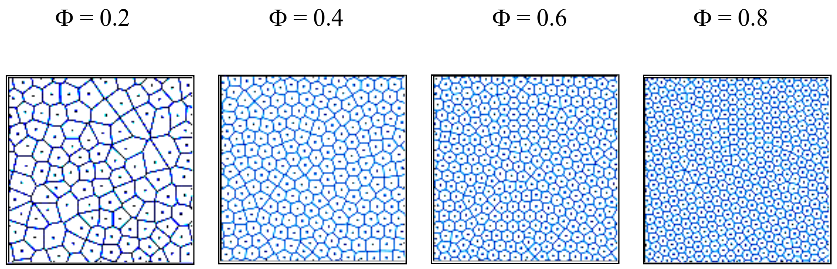


Figure S1: Representative Voronoi tessellations constructed from model particle centers, depicting the evolution of tissue morphology during quasistatic cellular densification

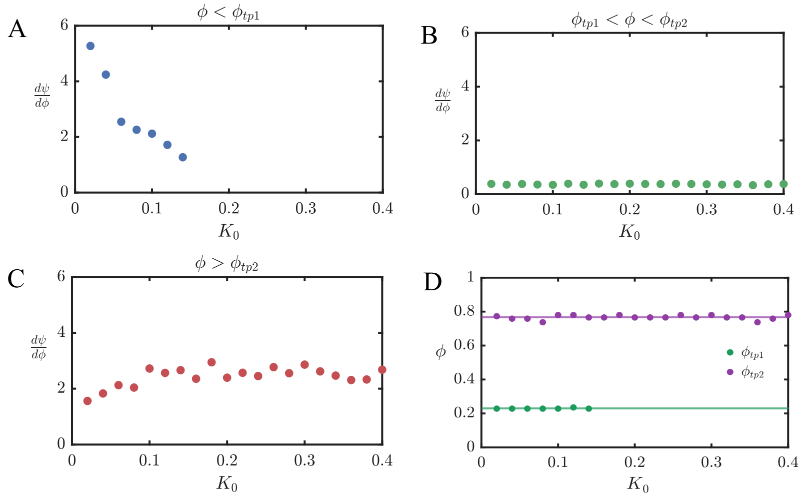


Figure S2: (A-C) Dependence of the ordering rate $d\psi/d\phi$ on the intrinsic division rate $K_0$ in the low-density regime ($\phi < \phi_{tp1}$), intermediate packing fraction regime ($\phi_{tp1} < \phi < \phi_{tp2}$),and in the high-density regime ($\phi > \phi_{tp2}$) respectively.(D) Transition packing fractions $\phi_{tp1}$ (green) and $\phi_{tp2}$ (purple) as functions of $K_0$. The green and purple line represents the average through all $K_0$.

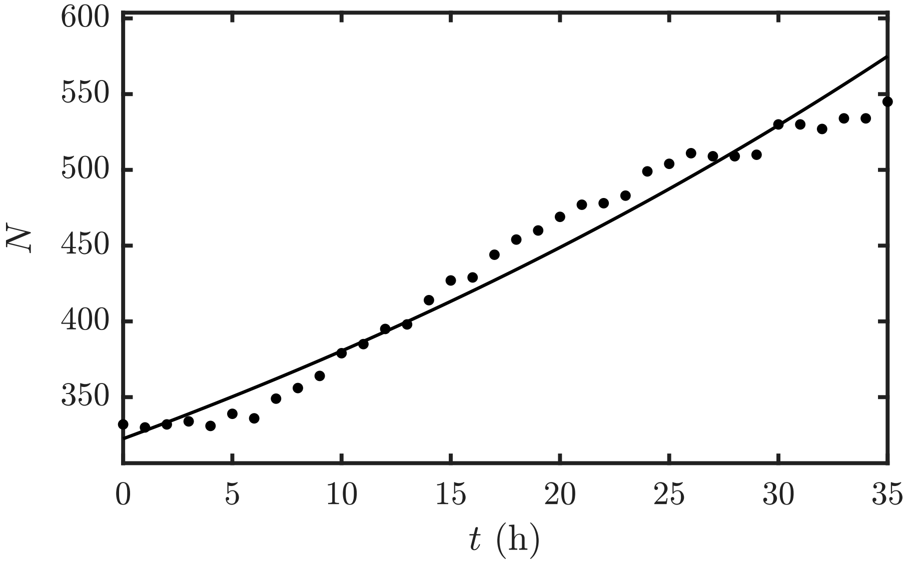


Figure S3: Experimentally observed change in cell number with time $N(t)$, where experimental data (scatter points) are overlaid with a spline fit(solid line).

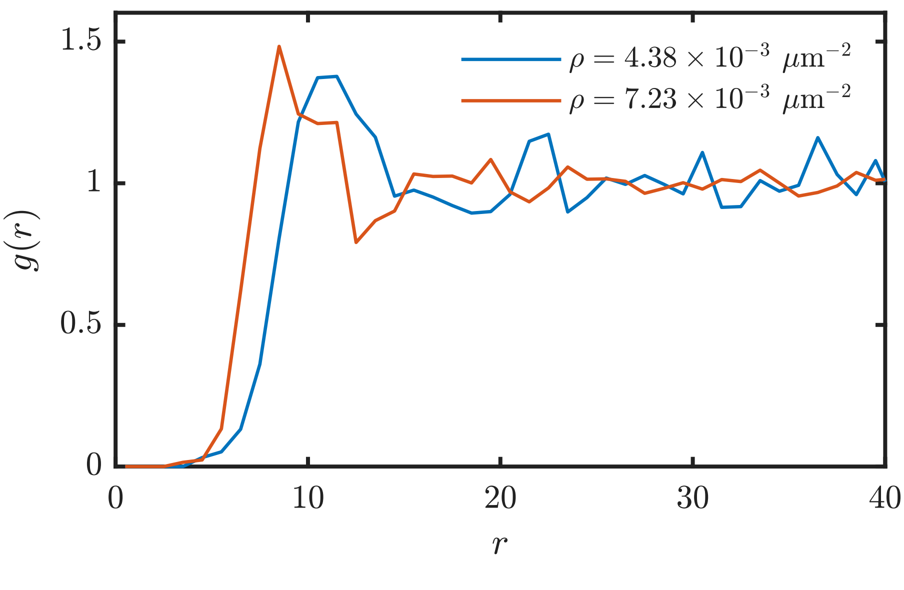


Figure S4: Radial distribution function $g(r)$ obtained from experimental snapshots from early (blue line, $\rho = 4.38 \times 10^{-3} \mu m^{-2}$) and late (red line, $\rho = 7.23 \times 10^{-3} \mu m^{-2}$) stage of densification respectively.
